# To FRET or Not to FRET: Bioinformatics and Fluorescence Spectroscopy suggest that Reduced Tryptophan–to–Heme Energy Transfer Facilitates Lignin Degradation in Class II Peroxidases

**DOI:** 10.64898/2025.12.31.697144

**Authors:** Yi Ren, Dale Ang, Haluk Ertan, Anne Poljak, Janice R. Aldrich-Wright, Wallace Bridge, Khawar Sohail Siddiqui

## Abstract

A key step in the evolution of lignin-degrading enzymes is revealed by the observation that, unlike other heme-proteins studied to date, Class II peroxidases exhibit minimal energy transfer from tryptophan (Trp) to heme residues. Bioinformatics analyses and molecular dynamics simulations of Class II (MnP and VP) and Class III (horseradish peroxidase, HrP) structures indicate that the Trp residue in HrP has the highest orientational factor and fluorescence resonance energy transfer (FRET) efficiency. By contrast, Trp residues in MnP and VP display low FRET efficiency due to unfavorable orientation factors despite their proximity to the heme. Steady-state fluorescence experiments confirmed this low FRET efficiency, showing strong emission in MnP and VP but weak emission in HrP. This decreased Trp-to-heme energy transfer appears to minimize competition between direct FRET and long-range electron transfer (LRET), allowing electrons to flow from bulky lignin substrates to the heme center. Such a mechanism likely provided a selective advantage during the evolution of Class II peroxidases, facilitating efficient lignin degradation at the enzyme surface.

**Highlights:** Class II peroxidases show reduced Trp-to-heme FRET compared with HRP

MD simulations reveal unfavorable Trp–heme orientation in VP and MnP

Low FRET correlates with strong Trp fluorescence in VP and MnP

Reduced FRET favors long-range electron transfer (LRET) during lignin oxidation

Findings suggest an evolutionary adaptation in lignin-degrading peroxidases

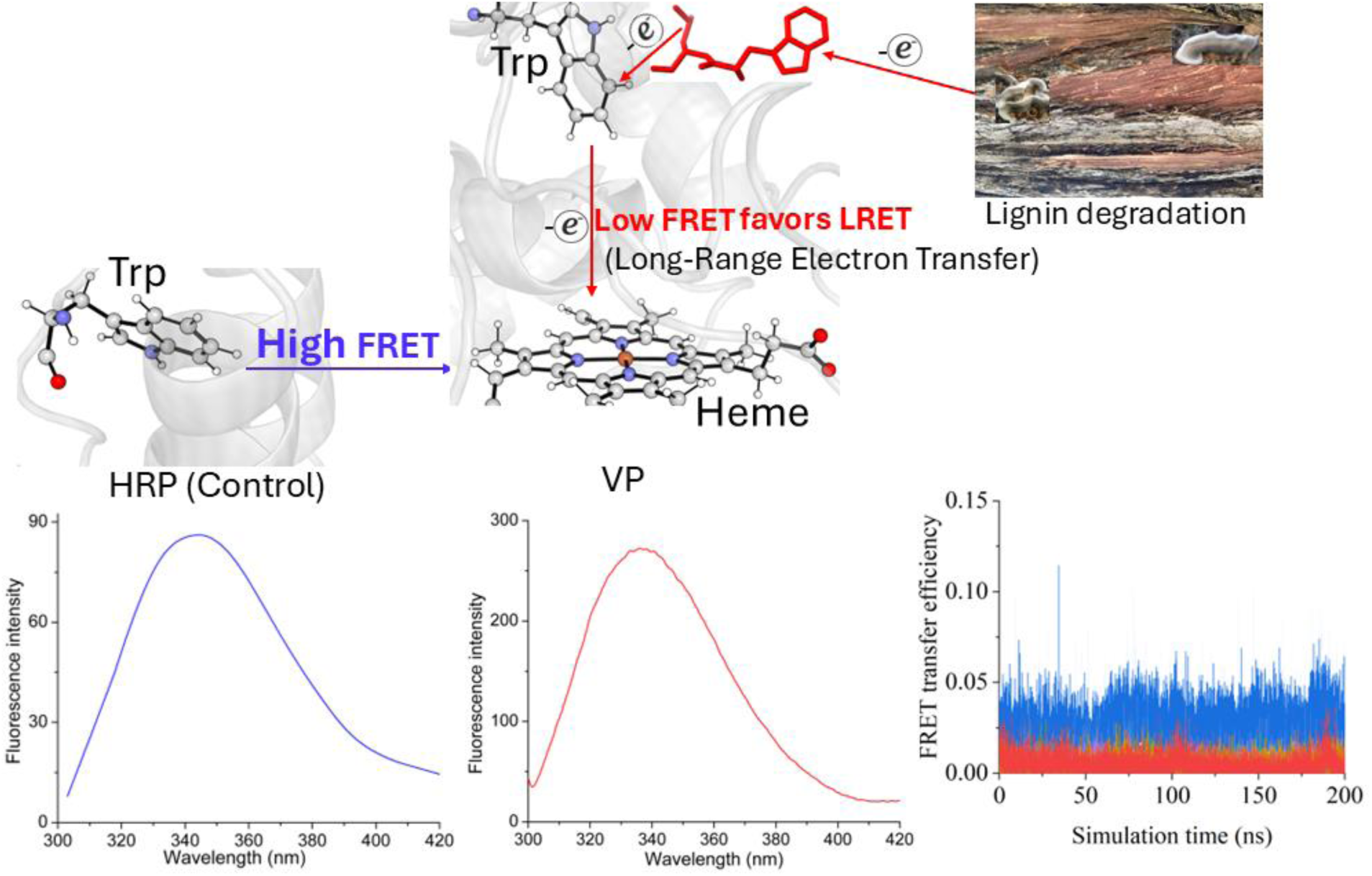

## 1. Introduction

Heme-peroxidases belong to the oxidoreductases (EC: 1.11.1.x) group of enzymes that catalyze the oxidation of a variety of substrates in the presence of H_2_O_2_. The non-animal heme-peroxidases found in bacteria, fungi, and plants are subdivided into three main classes (http://peroxibase.toulouse.inra.fr/). Class I consists of intracellular peroxidases such as yeast cytochrome c peroxidase (CcP). Class II consists of secretory fungal peroxidases such as lignin peroxidase (LiP), manganese peroxidase (MnP), and LiP-MnP hybrid versatile peroxidase (VP). Class III consists of secretory plant peroxidases such as horseradish peroxidase (HrP). The catalytic cycles of these heme-peroxidases are similar and consist of three main steps. Firstly, the O–O bond in H_2_O_2_ is cleaved by heme-Fe^+3^, resulting in the formation of compound I (Fe^+4^cation radical), which is then converted to compound II (Fe^+4^=O) (Ortmayer and Green, 2021).

Both compounds I and II act on a variety of electron donor substrates in two-electron oxidation/reduction reactions (Wong 2009, Ertan et al. 2012). It is noteworthy that both compounds have a high redox potential (ability to accept electrons) in Class II peroxidases (Banci et al. 1991, Wong 2009, Martinez 2007, Ertan et al. 2012), whereas Class I and III peroxidases have low redox potential (Mondal et al. 1996, Reinke 2001). This enables Class II peroxidases to oxidize more recalcitrant high-redox substrates such as those found in lignins and humic acids which provides them with diverse commercial applications across industries (Kersten et al. 1990, Martinez 2007, Wong 2009, Ertan 2012, Siddiqui et al. 2014).

In heme proteins studied to date, tryptophan fluorescence is quenched via Forster/fluorescence resonance energy transfer (FRET) to the heme group (Weber and Teale 1959, Henry and Hochstrasser 1987). The energy transfer from tryptophan residues depends on the distance and orientation relative to the heme (Cabral et al. 2002). The absence of Trp fluorescence quenching in apo-proteins lacking heme is taken as further evidence of energy transfer to the heme (Chattopadhyay and Mazumdar, 2000, Gryczynski et al. 1993). Yet to be investigated is the potential for quenching of tryptophan fluorescence by secretory fungal Class II peroxidases (lignin peroxidase, LiP; manganese peroxidase, MnP, and hybrid LiP-MnP versatile peroxidase, VP).

It has been shown that FRET can compete with electron transfer when the former is more efficient due to favorable orientation and distance between the Trp and heme (Monni et al. 2015). Computational and site-directed mutagenesis analysis has since shown that the buried Trp is also involved in the LRET pathway when electron flow from the lignin substrate→ surface Trp→buried Trp→heme (Acebes et al. 2017).

Here, we show for the first time that, unlike other heme proteins, Trp residues in MnP and VP do not exhibit fluorescence quenching. The distances and orientation angles between Trp residues and heme in Class II peroxidase *ab initio* structures, along with FRET efficiency calculated from molecular dynamics simulations (MDS), led to the hypothesis that energy transfer in VP and MnP may be very low. This hypothesis was experimentally tested using steady-state fluorescence, including spectral and unfolding analyses, on both MnP and VP, with HrP serving as a control. The minimal energy transfer observed in VP and MnP, and its implications for the evolution of Class II peroxidases and the degradation of the recalcitrant lignin substrates are discussed.

## 2. Materials and methods

### 2.1. Purification and modeling of peroxidases

Purified MnP from *Nematoloma (Phlebia) frowardii* and VP from *Bjerkendera adusta* were purchased from JenaBios, Germany and further purified using Mono-Q (HR 5/5) anion-exchange chromatography using 20 mM Tris/HCl, pH 7 as loading buffer and eluted with loading buffer + 1 M NaCl. The eluted proteins were simultaneously detected at dual wavelengths (280 and 405 nm). The fractions corresponding to a single 407 nm peak were collected and subjected to both MnP and LiP activity assays as described (Ertan et al. 2012). Fractions corresponding to the 407 nm heme peak and VP and MnP activities were pooled. The purified VP and MnP were subjected to 12% SDS-PAGE to assess their purity. Purified HrP was purchased from Sigma-Aldrich and was used without further purification.

High-resolution structures of HrP (9h1m) and VP from *Pleurotus eryngi*i (2boq) were downloaded from Protein Data Bank (https://www.rcsb.org/). *Ab initio* structures of VP from *Bjerkendera adusta* (Genbank: ABQ44529.1) and MnP from *Nematoloma frowardii* (*Phlebia sp*. B19 (GenBank: ABR66918.1) after removing the signal peptide were built using Alphafold 3 available at https://alphafoldserver.com/ (Abramson et al. 2024).

### 2.2. Molecular dynamic simulations (MDS)

The protein cofactor heme was parameterized at B3LYP/6-31G(d) level of theory (Becke, 1993, Frisch et al., 1984, Grimme, 2010, Krishnan et al., 1980, Stephens et al., 1994) using Gaussian 09 (Frisch et al., 2009). The charge of heme was set as 1 and spin multiplicity as 6 (Fe (III) and negative 2 for porphyrin ring). The Sobtop package developed by Lu was used to assign the generalized AMBER force field (GAFF) (Wang et al., 2004) atom types of the heme cofactor, the universal force field (UFF) atom type (Rappé et al., 1992) was applied for the ferric atom, while the remaining heme atoms were assigned GAFF atom types. This hybrid assignment approach, using UFF for transition metals and GAFF for organic atoms, has been previously adopted in similar studies (Keot and Sarma, 2024, Li et al., 2024). The atomic charges were calculated with Restrained Electrostatic Potential (RESP) scheme by using the Multiwfn wave function program (Lu and Chen, 2012).

Parameter validation of heme cofactor, including atomic names, charges, bonds, angles and dihedrals, is presented in the Supplementary Figures S1-S2 and Table S1. The validation results indicate the molecular mechanics parameters of heme cofactor are acceptable for subsequent molecular dynamics simulation. The predicted protein structures of VP_fold and MnP_fold were validated by aligning against the crystal structure of VP (PDB id: 2bop), Supplementary Figure S3.

All protein structures were parameterized using the Amber Software (Case et al. 2005) and ff14SB force field (Maier et al., 2015), and protonation states were assigned at pH 4.5 using CHARMM-GUI PDB Reader (Jo et al., 2008), and verified by PROPKA 3. Histidine residues were specified using AMBER residue names, including HID (Nδ-H), HIE (Nε-H), HIP (+1, both nitrogen atoms protonated). All systems were solvated in a cubic box of TIP3P water molecules (Jorgensen et al., 1983), ensuring a 10 Å buffer between the solute and the box edge.

All the simulations and the analysis of the trajectories were performed with GROMACS software version 2024.3-gpuvolta package (Abraham et al., 2015, Abraham et al., 2024). All the starting structures, PceA with ligand docked in the active site, were energy relaxed with 1000 steps of steepest-descent energy minimization. The systems were then equilibrated in two stages. Initially, a constant volume canonical ensemble (NVT) equilibration at 303.15k using V-rescale thermostat (Bussi et al., 2007) for 4 nanoseconds (ns) using a time step of 1 fs. This was followed by constant pressure isobaric-isothermal ensemble (NPT) equilibration, where the pressure was maintained at 1 bar by using Parrinello-Rahman barostat (Parrinello and Rahman, 1981, Ke et al., 2022), with an isotropic pressure coupling type. The NPT equilibration was carried out in 4 stages: position restraints were applied to the backbone and side chains of protein and cofactor, starting with force constants of 400 and 40 kJ/mol/nm², respectively, from the first stage of NVT equilibration, and gradually reduced by 100 and 10 kJ/mol/nm², respectively. At the last stage of equilibration, the system was fully relaxed. The Verlet cutoff scheme (Grubmüller et al., 1991) was employed, the cutoff radius for short-range van der Waals interactions was set to 1.2 nm. The long-range electrostatic interactions were treated using the Particle-Mesh Ewald (PME) method (Petersen, 1995). All bonds involving hydrogen atoms were constrained using the LINCS algorithm (Hess et al., 1997). After equilibration, three independent 200 ns production runs were launched using 2 fs time step. The SHAKE algorithm (Ryckaert et al., 1977) was used to constrain all bonds involving hydrogen atoms. All analyses were carried out using gmx rms, gmx angle, distance and gangle for vector angle measurements.

#### 2.2.1 FRET orientation factor computation

From the MD trajectories, the FRET orientation factor *k*^2^ was calculated as (Hunt et al., 2012):

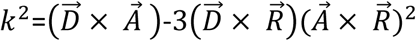

where 𝐷⃗ and 𝐴⃗ are the normalized transition dipole moment of the donor (tryptophan) and acceptor (HEM), respectively, (Stevens et al., 2012). 𝑅⃗ is the normalized distance vector between the donor and acceptor (Figure 1)

**Figure 1.**
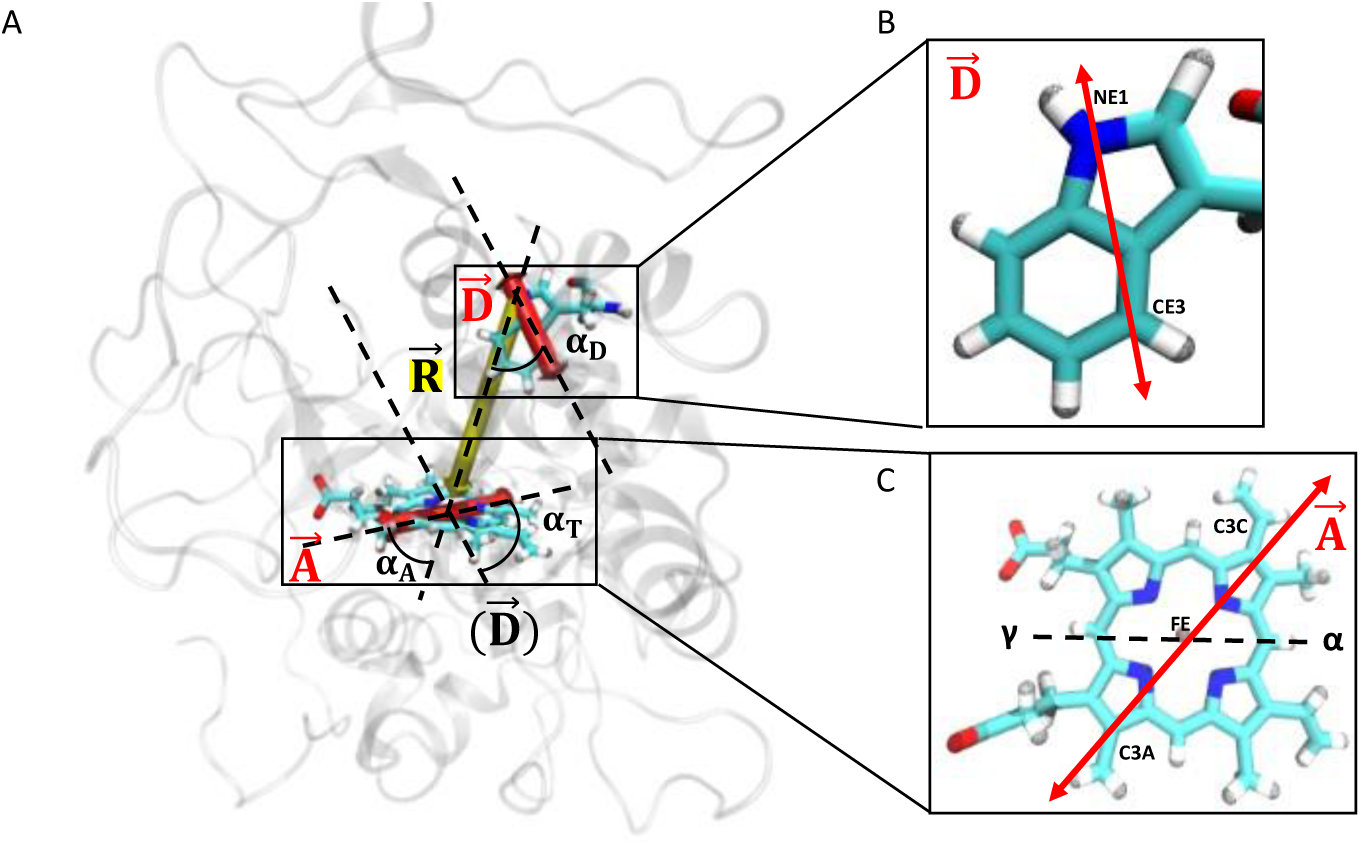
Illustration of the tryptophan and heme configuration (with VP as an example). (A) A representative VP donor and acceptor configuration. Donor (𝐷⃗ ) and acceptor (𝐴⃗) transition dipole vectors are shown in red. The distance vector 𝑅⃗ between the tryptophan and heme is shown in yellow. The angles between the vectors are 𝛼_𝐷_ (between 𝐷⃗ and 𝑅⃗ ), 𝛼_𝐴_ (between 𝐴⃗ and 𝑅⃗ ), and 𝛼_𝑇_ (between 𝐷⃗ and 𝐴⃗). (B) The tryptophan responsible for fluorescence emission is shown in the enlarged image, in which the atoms used for defining the Donor (𝐷⃗ ), represented as a red arrow and its direction is the vector that links atoms NE1-CE3 (red arrow). (C) The heme configuration as acceptor is shown in the enlarged image, in which the atoms used for defining the Acceptor (𝐴⃗), represented as a red arrow, and its direction is linked by atoms C3A-C3C.

The equation above can be rewritten as a function of the angle between vectors (Teijeiro-Gonzalez et al., 2021):

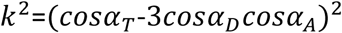

where 𝛼_𝑇_, 𝛼_𝐷_, and 𝛼_𝐴_ are the angles betwee^n⃗^𝐷⃗⃗ and 𝐴⃗, 𝐷⃗ and 𝑅⃗ , and 𝐴⃗ and 𝑅⃗ , respectively.

The FRET efficiency was calculated from the relative dipole orientation between the tryptophan and heme through the orientation factor *k*^2^ and their distance R with an approximation form (Van Beek et al., 2007):

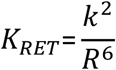

### 2.3. Fluorescence analysis

Spectral analysis and thermal unfolding of all peroxidases (0.20 mg mL^-1^) were monitored in a 0.1 M tartaric acid/NaOH, pH 4.5, buffer using a 0.2 mL capacity, 1 cm path length quartz cuvette and a PerkinElmer LS 50B spectrometer connected to a Peltier temperature controller. The protein solution was excited at 295 nm and a spectrum was taken from 302–450 nm at 4°C. The protein was then subjected to thermal unfolding from 4–94 °C at a ramp speed of 1.1 °C min^-1^ and monitored at an emission of 336 nm. After cooling to 4 °C, another spectrum was taken to determine the spectral intensity and shift due to unfolding. For all fluorescence experiments, excitation and emission slits were kept at 5 nm. All spectra were averaged from three scans at a scan rate of 25 nm/min.

## 3. Results

### 3.1. Molecular dynamic simulations (MDS)

AF3-predicted structures of VP-BA and MnP-Nf exhibited high confidence, with pTM scores of 0.97 and 0.96, respectively, and mean pLDDT scores exceeding 90%, indicating minimal expected positional errors. Structural superposition of AF3 models onto the high-resolution X-ray structure of VP from *Pleurotus eryngii* (2boq) revealed excellent alignment of the conserved Trp residues in both enzymes (Supplementary Figure S3), supporting the reliability of the AF3 models for VP-BA and MnP-Nf.

Hybrid VP comprises both manganese peroxidase (MnP) and lignin peroxidase (LiP) activities and has two Trp residues (Trp172 and Trp 252) as in LiP. *Ab initio* structures of VP and MnP superimposed on the X-ray structures of HrP (Figure 2) showed that the Trp 252 residue of VP has its indole side chain pointing towards the heme ring. This residue is homologous to Trp256 in MnP and is conserved. The second Trp residue in VP (Trp172) is located further away and its indole side chain points towards the protein surface, and away from the heme ring. This residue is unique to the long-range electron transfer (LRET) pathway in LiP and VP enzymes for the degradation of bulky lignin molecules at the enzyme surface, and is absent in MnP (Pogni et al. 2006, Wong 2009, Ertan 2012, Morales et al. 2012, Pérez-Boada et al. 2005).

**Figure 2.**
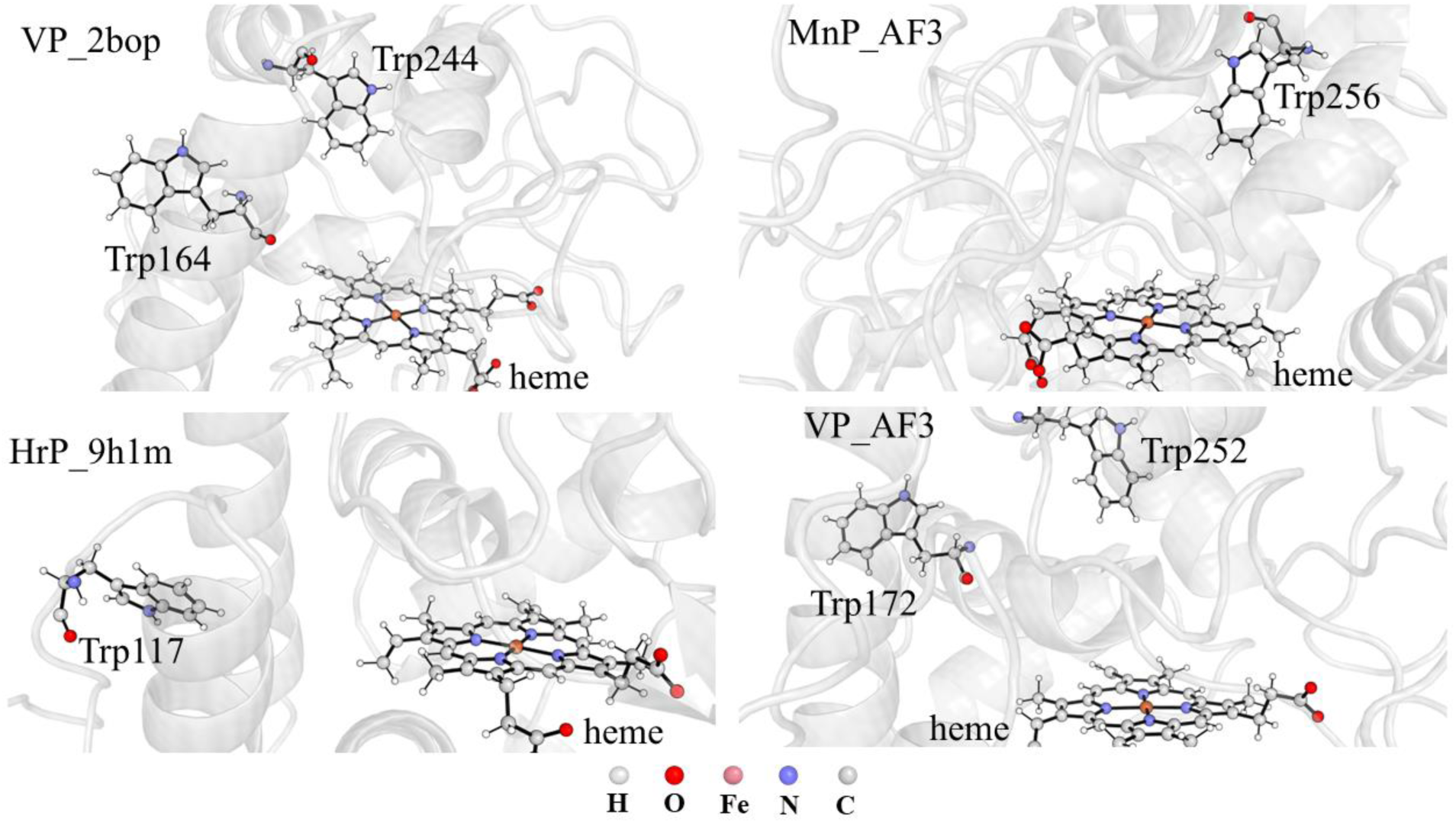
The aligned structure of HrP, VP, and MnP. The relative orientations of Heme and the key Trp residues of interest are shown as balls and sticks. MnP_AF3 and VP_AF3 structures are predicted by AlphaFold3.

At 200 ns, Trp117 in HrP exhibits the longest donor–acceptor distance but achieves the highest orientation factor and energy transfer efficiency due to its planar alignment with the heme. In contrast, Trp172 and Trp252 in VP and Trp256 in MnP show more favorable distances but lower orientation factors and energy transfer efficiencies due to their perpendicular configuration relative to the heme. These early trends highlight the importance of dipole alignment over distance in modulating energy transfer.

MDS at 200 ns revealed that the Trp–heme orientation factors in Class II peroxidases (VP and MnP) are less favorable for FRET-based energy transfer as comparing with HrP (pdb id: 9 h1 m), suggesting an absence of tryptophan fluorescence quenching. In contrast, HrP showed a favorable parallel alignment of Trp117 with the heme plane, resulting in the most efficient energy transfer and highest FRET orientation factor value.

To examine the FRET transfer efficiency, the approximated calculation was carried out (as shown in the Method Section 2.2.2) by using a 200 ns molecular dynamics simulation. To test the stability of the entire protein structures, the root mean square deviations of protein backbone atoms were measured. No significant fluctuation is exhibited over the 200 ns simulation, indicating protein structure stability (Figure 3A). Due to the significance of distance between the Trp residue and heme in the approximated FRET transfer efficiency, the distances between the Trp residues of each protein and heme were measured for the N atom and iron atoms of heme (Figure 3B) and did not exhibit significant variation across all the measurements across the 200 ns simulation time. As shown in Figure 3B, the Trp117 of HrP (blue), is separated by the greatest distance between the donor and acceptor, as compared with other Trp residues in VP and MnP.

**Figure 3.**
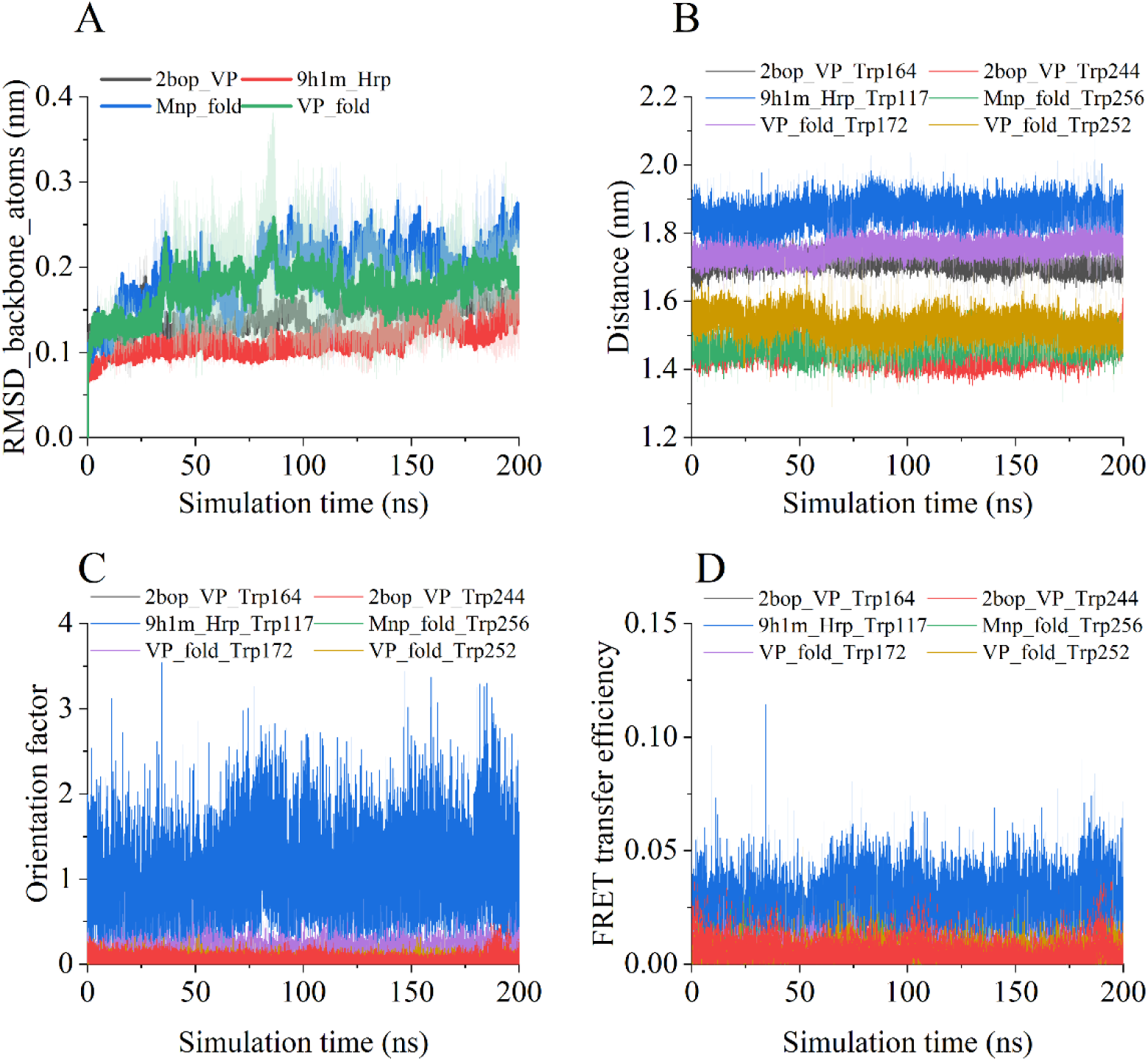
Molecular dynamics simulations for crystal structures of VP (PDB id: 2bop) and HrP (PDB id: 9h1m) and AlphFold predicted structures of MnP_fold and VP_fold. A) the Root Mean Square Deviation (RMSD) of protein backbone atoms over the 200 ns simulation time, showing the stability of protein structure. B) The distance showing the calculated Trp-to-heme energy transfer efficiency, measured between the N atom of the Trp residue and the iron of Heme cofactor. C) the Orientation factor. D) the FRET transfer efficiency from Trp to heme.

The approximated FRET transfer in the present study is the coupling between the Trp residues’ emission transition dipole and absorption transition dipole heme, which is subjected to the relative orientation between Trp residues and heme. Therefore, the orientation factor is calculated as per the approximated calculation in the method section 2.2. Across the 200 ns simulation, the angles 𝛼_𝑇_, 𝛼_𝐷_, and 𝛼_𝐴_ betwee^n⃗^𝐷⃗⃗ and 𝐴⃗, 𝐷⃗ and 𝑅⃗ , and 𝐴⃗ and 𝑅⃗ , respectively, are calculated for each frame. Trp117 of HrP (Figure 3C blue), has the highest variation compared to other Trp residues. Trp117 also has the highest orientation factor as compared with the other Trp residues in MnP and VP. This is because the configuration of Trp117 of HrP is within the plane of heme. The normalized transition dipole moment vectors of the donor (Trp) 𝐷⃗ and acceptor (HEM) 𝐴⃗ are in parallel positions, resulting in a higher orientation factor. By contrast, Trp172 (purple) and Trp252 (yellow) in VP, and Trp256 in MnP fold (green), have noticeably lower orientation factors due to their perpendicular position relative to heme. As for the calculated energy transfer efficiency (Figure 3D), Trp117 in HrP has the highest energy transfer efficiency, despite having the most unfavorable distance. Trp172 and Trp252 in VP and Trp256 in MnP have noticeably lower energy transfer efficiency, due to the unfavorable orientation factor, even though their distances are more favorable. To validate these computational findings, we next tested this prediction experimentally using steady-state fluorescence measurements to compare HrP (control) with VP and MnP.

### 3.2. Fluorescence Analysis Confirms the MDS-Based Hypothesis of Less Energy Transfer from Trp to Heme in Class II Peroxidases

Molecular dynamics simulation findings (Figure 3) suggested that, unlike HrP, Class II peroxidases might not be able to transfer the energy of Trp excitation to the heme. To test this hypothesis, we performed steady-state fluorescence experiments based on spectral analysis and thermal unfolding of VP, MnP, and HrP. Both VP (Supplementary Figure S4A) and MnP (result not shown) were subjected to anion-exchange chromatography. The SDS-PAGE showed a closely-spaced doublet band for VP (Supplementary Fig 4, inset) as described previously (Ertan et al. 2012) and a dark band for MnP (Supplementary Figure S4B).

Using equal concentrations of all three purified proteins (normalized by A_407_) under identical experimental conditions, the samples were subjected to steady-state fluorescence measurements (Figure 4). Fluorescence spectral analysis revealed that VP and MnP (Figure 4A, C, solid lines) exhibited significant fluorescence intensity at 4 °C when excited at 295 nm, with a maximum emission around 336 nm. By contrast, the HrP control displayed negligible intrinsic fluorescence, with a maximum emission at approximately 330 nm (Figure 4E, solid line). This indicates that energy transfer from Trp to the heme does not occur in VP and MnP, whereas in HrP, the Trp residue is quenched due to energy transfer from Trp117 to the heme.

**Figure 4.**
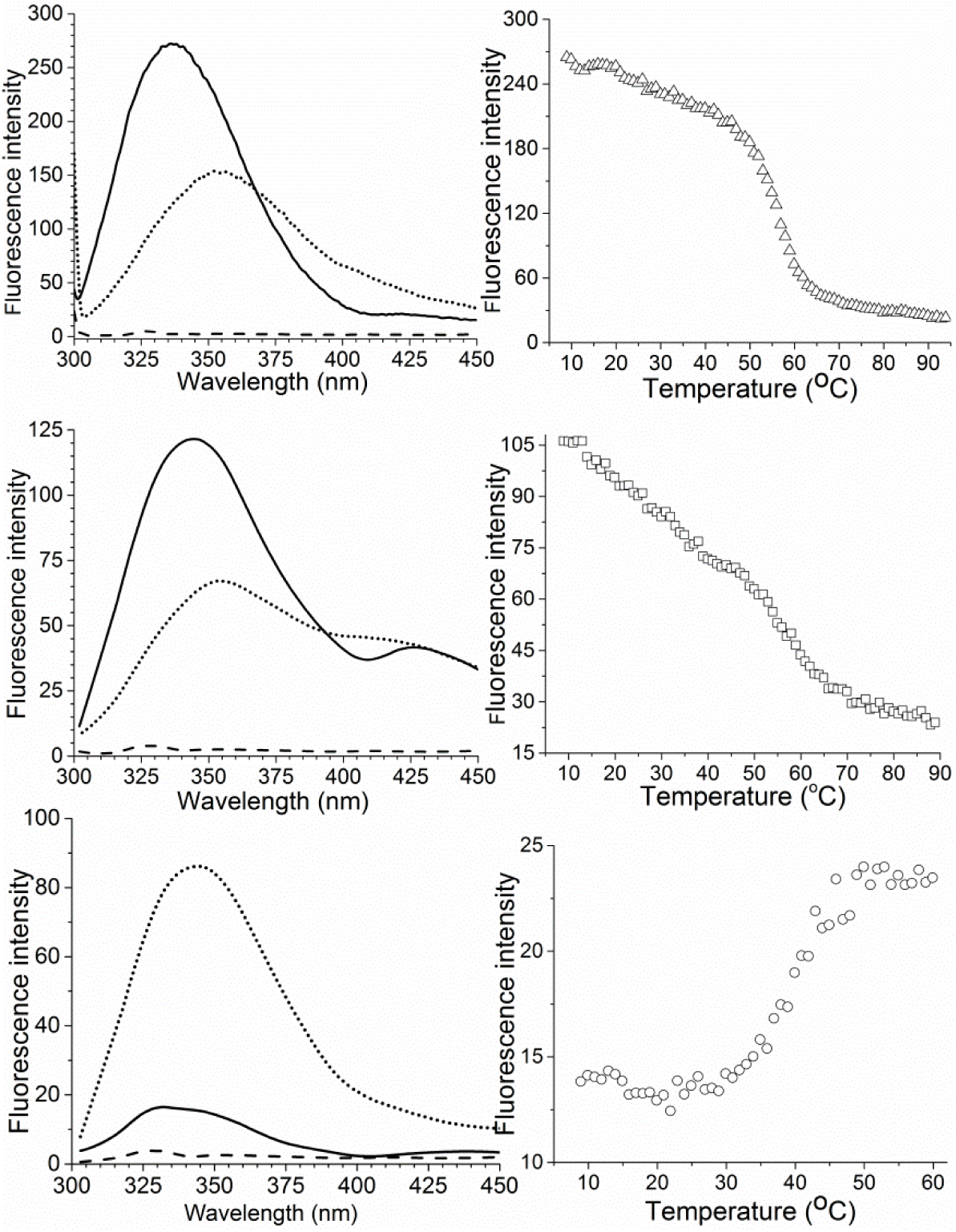
Fluorescence analysis of all three peroxidases. Left panels (A, C, E): spectrum analysis, and right panels (B, D, F): unfolding melting curves. Upper panels (A, B): VP, middle panels (C, D): MnP, and lower panels (E, F): HrP. Solid line: spectrum before unfolding at 8 °C, dotted line: spectrum at 8 °C after complete unfolding. Broken line: 0.1 M tartaric acid/NaOH, pH 4.5 buffer. Tryptophan residues were excited at 295 nm and the unfolding was monitored at 336 nm.

Furthermore, the red-shifted emission maxima of VP and MnP suggest that their Trp residues reside in a relatively less hydrophobic environment compared to HrP.

To further confirm that the Trp residue/s in VP and MnP do not transfer energy to heme, all three peroxidases were subjected to thermal unfolding and the fluorescence intensity was monitored at their respective emission maximum (Figure 4 BDF). The results demonstrated that MnP and VP exhibit a decrease in fluorescence intensity with progressive thermal unfolding due to exposure of the Trp residue/s to a more polar environment (Fig. 4BDF). By contrast, the thermal unfolding of HrP showed a progressive increase in fluorescence intensity with increased thermal unfolding (Figure 4F). This is interpreted as an increase in the Trp-heme distance and changes in their relative orientation, which decreases the Trp quenching as progressively less energy is transferred to the heme due to unfolding (Chattopadhyay and Mazumdar, 2000).

A second fluorescence scan after the complete unfolding of HrP (Figure 4E, dotted line) showed an increase in fluorescence intensity compared to that of unfolded HrP (Figure 4E, solid line) as well as red-shift of emission maximum to ∼350 nm thus confirming that changes in Trp-heme distance and orientation have resulted in reducing Trp quenching. On the other hand, the second fluorescence scan after the complete unfolding of VP (Figure 4A, dotted line) and MnP (Figure 4C, dotted line) showed a decrease in fluorescence intensity (Trp quenching) compared to that of unfolded VP and MnP (Figure 4AC, solid lines), implying exposure of hydrophobic residues to the more polar protein surface environment.

## 4. Discussion

We report for the first time that Class II peroxidases show low FRET efficiency from Trp to heme, unlike Class III peroxidases. This variation of structure and function of the ligninases (MnP, LiP, VP) now provides a mechanistic explanation for their ability to degrade bulky high-redox potential substrates such as lignin found in plant material. The removal of lignin from wood enables cellulolytic enzymes to further degrade exposed cellulose and hemicellulose (Guerriero et al. 2016).

All heme proteins, including non-animal peroxidases (Class I and III) studied to date show high FRET from Trp to heme, which depends on the relative Trp-heme distance and orientation (Weber and Teale 1959, Henry and Hochstrasser 1987, Chattopadhyay and Mazumdar, 2000, Cabral et al. 2002) and rotation of the heme around its α-γ-meso-axis (Gryczynski et al. 1993). This implies that when Trp residue/s are excited, the spectrum shows very little fluorescence emission intensity due to the transfer of energy from Trp to heme. The fluorescence intensity increases because of protein unfolding and the resulting change in the Trp to heme distances and orientations, as shown in the case of Class I (such as cytochrome c) and Class III (such as HrP) model peroxidases (Tsaprailis et al. 1998, Chattopadhyay and Mazumdar, 2000). Conversely, in apo-proteins lacking heme, Trp fluorescence quenching does not occur (Gryczynski et al. 1993).

The interpretation of the MD simulation results can reveal both the orientation and distance fluctuations that influence the FRET efficiencies between Trp and heme. The FRET efficiencies in the tested peroxidases are highly sensitive to the heme-Trp separation, which decreases with the inverse 6^th^ power of the donor-acceptor distance (refer to the formula above). In addition, FRET efficiencies are also highly dependent on the mutual orientation factor between the donor-acceptor transition dipoles (De Torres et al., 2016), which is calculated from their angles.

Based on the Trp-heme angle, a parallel orientation (within the plane) facilitates FRET efficiency, whereas a perpendicular Trp-heme orientation (out of the plane) hinders energy transfer (Gryczynski et al. 1993, Cabral et al. 2002). In accordance with the present study, the calculated orientation factor for Trp at the parallel position with heme is noticeably higher than the Trp at the perpendicular position (Stevens et al., 2012). As previously reported, the inefficiency in quenching of nucleotide donor fluorescence is in accordance with the near-zero orientation factor (Lewis et al., 2005). Even with a long-distance separation between donor and acceptor, the FRET efficiency is more favorable due to the higher values of orientation factors (Rindermann et al., 2011). Therefore, the dependence of the orientation factor on FRET efficiency supports the possibility that the perpendicular orientation between donor-acceptor impedes the energy transfer.

Based on the distances and orientations of both Trp residues in VP and MnP, we expected these proteins to show significant fluorescence emission due to a lack of energy transfer to the heme. We subjected all three heme proteins to steady-state fluorescence spectral analysis and thermal unfolding. We employed HrP as a control in which its lone Trp (Trp117) residue (Figure 2) is known to be quenched by energy transfer to the heme (Tsaprailis et al. 1998,

Chattopadhyay and Mazumdar, 2000). There is no corresponding HrP Trp residue at the equivalent position in either VP or MnP (Figure 2). MnP was used as an additional example as it includes the conserved Trp (Trp256) residue located at an equivalent position in VP (which has its side chain pointing towards the heme ring) but lacks the LRET pathway surface Trp residue (Figure 2).

The experimental data for VP and MnP (Figure 4ABCD) contrast with the data for HrP (Figure 4EF), confirming the hypothesis based on MDS data that in VP and MnP minimal energy is transferred from Trp to the heme moiety. This results in an increased fluorescence intensity (before thermal unfolding) due to the unfavorable orientation factor.

Under UV excitation, Trp residues need to relax from high energy back to the ground state. There are two mechanisms for energy relaxation (fluorescence quenching) of Trp residues, the FRET and electron transfer, which compete (Monni et al., 2015, Consani et al., 2013, Glandières et al., 2007). Electron transfer can dominate when FRET is weak due to unfavorable distance and orientation factors (Monni et al., 2015). VP and MnP showed weak FRET efficiency, which would result in Trp relaxation through electron transfer to heme.

The capacity for the buried Trp residue with very low FRET to transfer electrons to heme has catalytic significance by forming a long-range electron transfer (LRET) pathway (Figure 2). The LRET pathway provides a means by which the bulky lignin polymer can be degraded by VP without the substrate coming in direct contact with the heme (Acebes et al., 2017). The electron is initially abstracted by the surface Trp (172 in VP), transferring it to the buried Trp (252 in VP) via a neighboring Phe. The role of the surface and buried Trp in the LRET is mapped by quantum mechanical/molecular mechanical (QM/MM) computational analysis and verified experimentally using site-directed mutagenesis (Acebes et al. 2017). Protein engineering experiments have shown that any mutation of Trp residues (surface 172 and/or buried 252) will compromise the lignin-catalyzing activity (Acebes et al., 2017). By contrast, MnP, which has a single Trp256 with near-zero FRET and without the surface Trp, is unable to degrade lignin.

Conversely, MnP has been converted into VP by introducing the surface Trp (Timofeevski et al., 1999), thereby activating the LRET pathway. This shows that MnP due to near-zero FRET has the necessary pathway for the transfer of electrons via the buried Trp to heme (Figure 2). The only difference is that instead of Phe in VP, there is an equivalent Leu between the two Trp residues (Morgenstern et al. 2008).

We speculate that such unique features of MnP and VP (Class 2 peroxidases) could have evolved 280 million years ago during the Permian era, when the huge amount of UV-B radiation (280–315 nm) exposure due to absence of the ozone layer may have contributed to mutations, and origin of new functions in the surviving species after the mass extinction (Visscher et al., 2004). At approximately the same time lignin-degrading fungi evolved, with the identification of a sharp decline in the rate of organic carbon burial. This marked the beginning of lignin degradation as a huge amount of wood lignin was recycled by Class II peroxidases (Floudas et al., 2012). The high intensity of UV-B in the Permian period is in the same range as that required for the excitation of Trp residues. Due to favorable FRET, most peroxidases were unable to transfer an electron from the buried Trp to heme and as a result, lacked a functional LRET pathway to degrade C-C and C-O-C bonds in lignin. By contrast, mutations that positioned Trp residues in unfavorable orientations for FRET minimized energy transfer, potentially enabled electron transfer from Trp to heme instead. This may have contributed to the emergence of a long-range electron transfer (LRET) pathway, in which electrons flow from surface-exposed Trp to the heme via an internal Trp, facilitating the degradation of bulky and recalcitrant lignin.

Notably, the evolutionary refinement of Class II peroxidases from ancestral forms is marked by a progressive decrease in the number of Trp residues—culminating in a single Trp in MnP—likely to optimize and regulate electron flow (Shahid et al. 2021).

## Conclusion

This study uncovers a key mechanistic distinction in the evolution of lignin-degrading peroxidases. Through integrated computational and experimental analyses, we demonstrate that Class II peroxidases (VP and MnP) lack enough fluorescence resonance energy transfer (FRET) from tryptophan to heme due to unfavorable orientation factors—unlike HrP, where FRET quenches Trp fluorescence efficiently. Instead, VP and MnP exploit a long-range electron transfer (LRET) pathway, enabling electrons to travel from surface-accessible Trp residues to the deeply buried heme center. This mechanism allows these enzymes to catalyze lignin degradation at a distance, overcoming steric barriers imposed by the bulky polymer.

Our findings provide a mechanistic basis for how structural adaptations in Class II peroxidases may have evolved to enable the efficient breakdown of lignin during a pivotal stage in fungal and enzymatic evolution. Future work should experimentally dissect the interplay between FRET and electron transfer in Class II peroxidases.

## Acknowledgements

This work was supported by the Australian Research Council. We acknowledge CHATGPT for editing English, word management, and formatting references.

## Author contributions

K.S.S., and H.E. conceived the original idea. K.S.S., H.E., Y.R, and D.A. designed the study. K.S.S. H.E., D.A., and Y.R performed the study. K.S.S., H.E., A.P., Y.R., D.A., W.B. Y.R., and J.R.A. discussed the data and wrote or edited the paper. All authors approved the final version of the manuscript.

## References

1. Abramson J, Adler J, Dunger J, et al. Accurate structure prediction of biomolecular interactions with AlphaFold 3. Nature. 2024;630:493–500. doi:10.1038/s41586-024-07487-w.

2. Abraham M, Alekseenko A, Basov V, Bergh C, Briand E, Brown A, Doijade M, Fiorin G, Fleischmann S, Gorelov S, et al. GROMACS 2024.3 Manual. 2024.

3. Abraham MJ, Murtola T, Schulz R, Páll S, Smith JC, Hess B, Lindahl E. GROMACS: high performance molecular simulations through multi-level parallelism from laptops to supercomputers. SoftwareX. 2015;1–2:19–25. doi:10.1016/j.softx.2015.06.001.

4. Acebes S, Ruiz-Dueñas FJ, Toubes M, Saez-Jimenez V, Perez-Boada M, Lucas MF, Martínez AT, Guallar V. Mapping the long-range electron transfer route in ligninolytic peroxidases. J Phys Chem B. 2017;121:3946–3954. doi:10.1021/acs.jpcb.7b01634.

5. Banci L, Bertini I, Turano P, Tien M, Kirk TK. Proton NMR investigation into the basis for the relatively high redox potential of lignin peroxidase. Proc Natl Acad Sci U S A. 1991;88:6956–6960. doi:10.1073/pnas.88.16.6956.

6. Becke AD. Density-functional thermochemistry. III. The role of exact exchange. J Chem Phys. 1993;98:5648–5652. doi:10.1063/1.464913.

7. Bussi G, Donadio D, Parrinello M. Canonical sampling through velocity rescaling. J Chem Phys. 2007;126:014101. doi:10.1063/1.2408420.

8. Cabral CB, Imasato H, Rosa JC, Laure HJ, da Silva CHTP, Tabak M, Garratt RC, Greene LJ. Fluorescence properties of tryptophan residues in the monomeric d-chain of *Glossoscolex paulistus* hemoglobin: an interpretation based on a comparative molecular model. Biophys Chem. 2002;97:139–157. doi:10.1016/S0301-4622(02)00046-7.

9. Case DA, Cheatham TE III, Darden T, Gohlke H, Luo R, Merz KM Jr, Onufriev A, Simmerling C, Wang B, Woods RJ. The Amber biomolecular simulation programs. J Comput Chem. 2005;26:1668–1688. doi:10.1002/jcc.20290.

10. Chattopadhyay K, Mazumdar S. Structural and conformational stability of horse-radish peroxidase: effect of temperature and pH. Biochemistry. 2000;39:263–270. doi:10.1021/bi990729o.

11. Consani C, Auböck G, van Mourik F, Chergui M. Ultrafast tryptophan-to-heme electron transfer in myoglobins revealed by UV 2D spectroscopy. Science. 2013;339:1586–1589. doi:10.1126/science.1231161.

12. De Torres J, Mivelle M, Moparthi SB, Rigneault H, van Hulst NF, Garcia-Parajo MF, Margeat E, Wenger J. Plasmonic nanoantennas enable forbidden Förster dipole–dipole energy transfer and enhance FRET efficiency. Nano Lett. 2016;16:6222–6230. doi:10.1021/acs.nanolett.6b02636.

13. Ertan H, Siddiqui KS, Muenchhoff J, Charlton T, Cavicchioli R. Kinetic and thermodynamic characterization of the functional properties of a hybrid versatile peroxidase using isothermal titration calorimetry: insight into manganese peroxidase activation and lignin peroxidase inhibition. Biochimie. 2012;94:1221–1231. doi:10.1016/j.biochi.2012.02.012.

14. Floudas D, Binder M, Riley R, Barry K, Blanchette RA, Henrissat B, Martínez AT, Otillar R, Spatafora JW, Yadav JS. The Paleozoic origin of enzymatic lignin decomposition reconstructed from 31 fungal genomes. Science. 2012;336:1715–1719. doi:10.1126/science.1221748.

15. Frisch MJ, Pople JA, Binkley JS. Self-consistent molecular orbital methods. 25. Supplementary functions for Gaussian basis sets. J Chem Phys. 1984;80:3265–3269. doi:10.1063/1.447079.

16. Frisch MJ, Trucks GW, Schlegel HB, et al. Gaussian 09 Revision D.01. Wallingford, CT: Gaussian Inc.; 2009.

17. Glandières J-M, Twist C, Haouz A, Zentz C, Alpert B. Resolved fluorescence of the two tryptophan residues in horse apomyoglobin. Photochem Photobiol. 2007;71:382–386. doi:10.1562/0031-8655(2000)071<0382:RFOTTW>2.0.CO;2.

18. Grimme S. DFT-D3 – a dispersion correction for density functionals, Hartree–Fock and semi-empirical quantum chemical methods. J Chem Phys. 2010;132:154104. doi:10.1063/1.3382344.

19. Grubmüller H, Heller H, Windemuth A, Schulten K. Generalized Verlet algorithm for efficient molecular dynamics simulations with long-range interactions. Mol Simul. 1991;6:121–142. doi:10.1080/08927029108022457.

20. Gryczynski Z, Fronticelli C, Tenenholz T, Bucci E. Effect of disordered hemes on energy transfer rates between tryptophans and heme in myoglobin. Biophys J. 1993;65:1951–1958. doi:10.1016/S0006-3495(93)81266-9.

21. Guerriero G, Hausman J-F, Strauss J, Ertan H, Siddiqui KS. Lignocellulosic biomass: biosynthesis, degradation, and industrial utilization. Eng Life Sci. 2016;16:1–16. doi:10.1002/elsc.201400196.

22. Henry ER, Hochstrasser RM. Molecular dynamics simulations of fluorescence polarization of tryptophans in myoglobin. Proc Natl Acad Sci U S A. 1987;84:6142–6146. doi:10.1073/pnas.84.17.6142.

23. Hess B, Bekker H, Berendsen HJ, Fraaije JG. LINCS: a linear constraint solver for molecular simulations. J Comput Chem. 1997;18:1463–1472. doi:10.1002/(SICI)1096-987X(199709)18:12<1463::AID-JCC4>3.0.CO;2-H.

24. Hunt J, Keeble AH, Dale RE, Corbett MK, Beavil RL, Levitt J, Swann MJ, Suhling K, Ameer-Beg S, Sutton BJ, Beavil AJ. A fluorescent biosensor reveals conformational changes in human immunoglobulin E Fc. J Biol Chem. 2012;287:17459–17470. doi:10.1074/jbc.M112.347559.

25. Jo S, Kim T, Iyer VG, Im W. CHARMM-GUI: a web-based graphical user interface for CHARMM. J Comput Chem. 2008;29:1859–1865. doi:10.1002/jcc.20945.

26. Jorgensen WL, Chandrasekhar J, Madura JD, Impey RW, Klein ML. Refined TIP3P model for water. J Chem Phys. 1983;79:926–935. doi:10.1063/1.445869.

27. Ke Q, Gong X, Liao S, Duan C, Li L. Effects of thermostats/barostats on physical properties of liquids by molecular dynamics simulations. J Mol Liq. 2022;365:120030. doi:10.1016/j.molliq.2022.120030.

28. Keot N, Sarma M. Unraveling the stability and magnetic properties of bis-hydrated Mn(II) complexes via tailored ligand design. J Phys Chem A. 2024;128:8346–8359. doi:10.1021/acs.jpca.4c03053.

29. Kersten PJ, Kalyanaraman B, Hammel KE, Reinhammar B, Kirk TK. Comparison of lignin peroxidase, horseradish peroxidase and laccase in the oxidation of methoxybenzenes. Biochem J. 1990;268:475–480. doi:10.1042/bj2680475.

30. Krishnan R, Binkley JS, Seeger R, Pople JA. Self-consistent molecular orbital methods. XX. A basis set for correlated wave functions. J Chem Phys. 1980;72:650–654. doi:10.1063/1.438955.

31. Lewis FD, Zhang L, Zuo X. Orientation control of fluorescence resonance energy transfer using DNA as a helical scaffold. J Am Chem Soc. 2005;127:10002–10003. doi:10.1021/ja052054k.

32. Li Y, Jin X, Moubarak E, Smit B. A refined set of universal force field parameters for some metal nodes in metal–organic frameworks. J Chem Theory Comput. 2024;20:10540–10552. doi:10.1021/acs.jctc.4c01113.

33. Lu T. SOBTOP Version 1.0. Available at: http://sobereva.com/soft_sobtop

34. Lu T, Chen F. Multiwfn: a multifunctional wavefunction analyzer. J Comput Chem. 2012;33:580–592. doi:10.1002/jcc.22885.

35. Maier JA, Martinez C, Kasavajhala K, Wickstrom L, Hauser KE, Simmerling C. ff14SB: improving the accuracy of protein side chain and backbone parameters from ff99SB. J Chem Theory Comput. 2015;11:3696–3713. doi:10.1021/acs.jctc.5b00255.

36. Martinez AT. High redox potential peroxidases. In: Polaina J, MacCabe AP, editors. Industrial enzymes. Springer; 2007. p. 475–486. doi:10.1007/1-4020-5377-0_27.

37. Mondal MS, Fuller HA, Fraser A. Direct measurement of the reduction potential of catalytically active cytochrome c peroxidase compound I. J Am Chem Soc. 1996;118:263–264. doi:10.1021/ja953303j.

38. Morales M, Mate MJ, Romero A, Martínez MJ, Martínez ÁT, Ruiz-Dueñas FJ. Two oxidation sites for low redox potential substrates. J Biol Chem. 2012;287:41053–41067. doi:10.1074/jbc.M112.405548.

39. Morgenstern I, Klopman S, Hibbett DS. Molecular evolution and diversity of lignin-degrading heme peroxidases in the Agaricomycetes. J Mol Evol. 2008;66:243–257. doi:10.1007/s00239-008-9086-x.

40. Monni R, Al Haddad A, van Mourik F, Auböck G, Chergui M. Tryptophan-to-heme electron transfer in ferrous myoglobins. Proc Natl Acad Sci U S A. 2015;112:5602–5606. doi:10.1073/pnas.1502731112.

41. Ortmayer M, Green AP. Heme peroxidases. In: Roberts GCK, Watts A, editors. Encyclopedia of Biophysics. Springer; 2021. doi:10.1007/978-3-642-35943-9_185-1.

42. Parrinello M, Rahman A. Polymorphic transitions in single crystals: a new molecular dynamics method. J Appl Phys. 1981;52:7182–7190. doi:10.1063/1.328693.

43. Pérez-Boada M, Ruiz-Dueñas FJ, Pogni R, Basosi R, Choinowski T, Martínez MJ, Piontek K, Martínez AT. Versatile peroxidase oxidation of high redox potential aromatic compounds. J Mol Biol. 2005;354:385–402. doi:10.1016/j.jmb.2005.09.047.

44. Petersen HG. Accuracy and efficiency of the particle mesh Ewald method. J Chem Phys. 1995;103:3668–3679. doi:10.1063/1.470043.

45. Pogni R, Baratto MC, Teutloff C, Giansanti S, Ruiz-Dueñas FJ, Choinowski T, Piontek K, Martínez AT, Lendzian F, Basosi R. A tryptophan neutral radical in the oxidized state of versatile peroxidase. J Biol Chem. 2006;281:9517–9526. doi:10.1074/jbc.M512070200.

46. Rappé AK, Casewit CJ, Colwell K, Goddard WA III, Skiff WM. UFF: a full periodic table force field for molecular mechanics and molecular dynamics simulations. J Am Chem Soc. 1992;114:10024–10035. doi:10.1021/ja00051a040.

47. Reineke W. Aerobic and anaerobic biodegradation potentials of microorganisms. In: Beek B, editor. Biodegradation and Persistence. Springer; 2001. p. 1–161. doi:10.1007/10508767_1.

48. Rindermann JJ, Akhtman Y, Richardson J, Brown T, Lagoudakis PG. Gauging the flexibility of fluorescent markers for the interpretation of fluorescence resonance energy transfer. J Am Chem Soc. 2011;133:279–285. doi:10.1021/ja108411k.

49. Ryckaert J-P, Ciccotti G, Berendsen HJ. Numerical integration of the Cartesian equations of motion of a system with constraints. J Comput Phys. 1977;23:327–341. doi:10.1016/0021-9991(77)90098-5.

50. Shahid M, Manoharadas S, Altaf M, Alrefaei AF. Organochlorine pesticides negatively influenced Enterobacter cloacae strain EAM 35. ACS Omega. 2021;6:5548–5559. doi:10.1021/acsomega.0c05856.

51. Siddiqui KS, Ertan H, Charlton T, Poljak A, Daud Khaled AK, Yang X, Marshall G, Cavicchioli R. Versatile peroxidase degradation of humic substances. J Biotechnol. 2014;178:1–11. doi:10.1016/j.jbiotec.2014.02.012.

52. Stephens PJ, Devlin FJ, Chabalowski CF, Frisch MJ. Ab initio calculation of vibrational absorption and circular dichroism spectra. J Phys Chem. 1994;98:11623–11627. doi:10.1021/j100096a001.

53. Stevens JA, Link JJ, Zang C, Wang L, Zhong D. Ultrafast dynamics of nonequilibrium resonance energy transfer in myoglobin. J Phys Chem A. 2012;116:2610–2619. doi:10.1021/jp211075g.

54. Teijeiro-Gonzalez Y, Crnjar A, Beavil AJ, Beavil RL, Nedbal J, Le Marois A, Molteni C, Suhling K. Time-resolved fluorescence anisotropy and molecular dynamics analysis of a novel GFP homo-FRET dimer. Biophys J. 2021;120:254–269. doi:10.1016/j.bpj.2020.11.2265.

55. Timofeevski SL, Nie G, Reading NS, Aust SD. Addition of veratryl alcohol oxidase activity to manganese peroxidase. Biochem Biophys Res Commun. 1999;256:500–504. doi:10.1006/bbrc.1999.0364.

56. Tsaprailis G, Chan DW, English AM. Conformational states in denaturants of cytochrome c and horseradish peroxidases. Biochemistry. 1998;37:2004–2016. doi:10.1021/bi971032a.

57. Van Beek DB, Zwier MC, Shorb JM, Krueger BP. Fretting about FRET: correlation between κ² and R. Biophys J. 2007;92:4168–4178. doi:10.1529/biophysj.106.100198.

58. Visscher H, van Looy CV, Collinson ME, Brinkhuis H, van Konijnenburg-van Cittert JH, Kürschner WM, Sephton MA. Environmental mutagenesis during the end-Permian ecological crisis. Proc Natl Acad Sci U S A. 2004;101:12952–12956. doi:10.1073/pnas.0404472101.

59. Wang J, Wolf RM, Caldwell JW, Kollman PA, Case DA. Development and testing of a general AMBER force field. J Comput Chem. 2004;25:1157–1174. doi:10.1002/jcc.20035.

60. Weber G, Teale FJW. Electronic energy transfer in haem proteins. Discuss Faraday Soc. 1959;27:134–141. doi:10.1039/DF9592700134.

61. Wong DWS. Structure and action mechanism of ligninolytic enzymes. Appl Biochem Biotechnol. 2009;157:174–209. doi:10.1007/s12010-008-8279-z.

