## Supplemental Material for "To FRET or Not to FRET: Bioinformatics and Fluorescence Spectroscopy suggest that Reduced Tryptophan–to–Heme Energy Transfer Facilitates Lignin Degradation in Class II Peroxidases"

#### Validation of the MD parameters for the heme cofactor

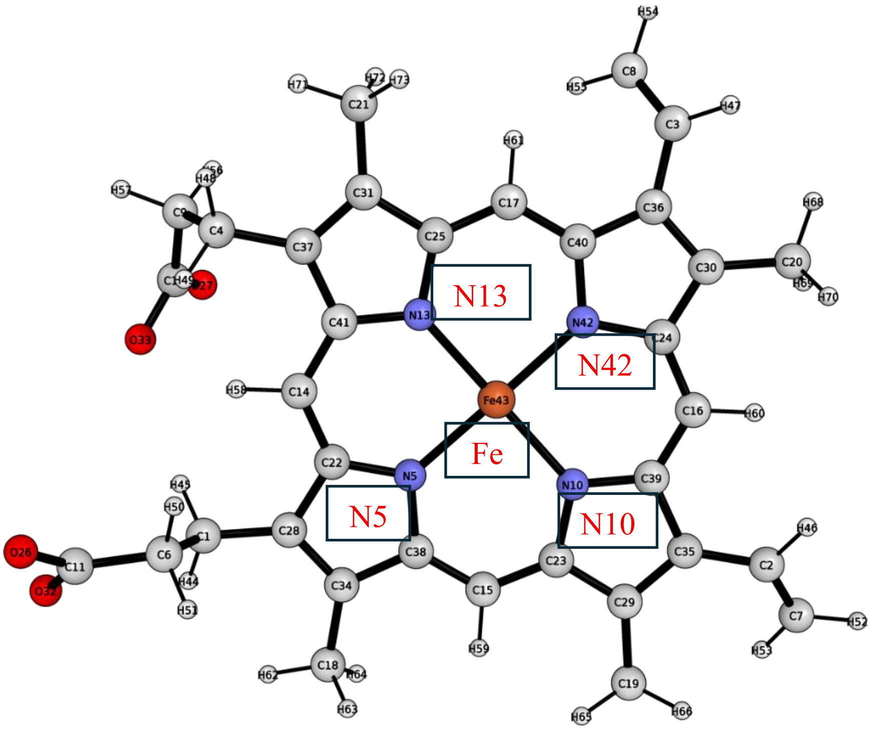

**Supplementary Figure 1.** A selected view of the heme cofactor and atoms used to validate the structure in the MD simulation.

**Supplementary Table 1.** Optimised structure measurements of the heme cofactor, calculated at B3LYP/ 6-31G(d) level of theory using Gaussian 09. The bonds and angles were used to validate the geometry in the MD simulation.

| Optimized structure of Heme and nitrogen atom | |
|---|---|
| N13_Fe_bond_distance (Å) | 1.96238 |
| N42_Fe_bond_distance (Å) | 1.95996 |
| N10_ Fe _bond_distance (Å) | 1.96077 |
| N5_ Fe _bond_distance (Å) | 1.95579 |
| angle_N13_Fe_N42 (Degree °) | 90.10468 |
| angle_N42_ Fe _N10 (Degree °) | 90.20896 |
| angle_N10_ Fe _N5 (Degree °) | 90.14635 |
| angle_N5_ Fe _N13 (Degree °) | 90.05234 |

For validating the molecular mechanics parameters of the heme cofactor, the distances between the N42 and Fe and N5 and Fe, angle of N13-Fe-N42 and N5-Fe-N10 were selected to measure the stability and compared with the parameters from the geometry optimised structure.

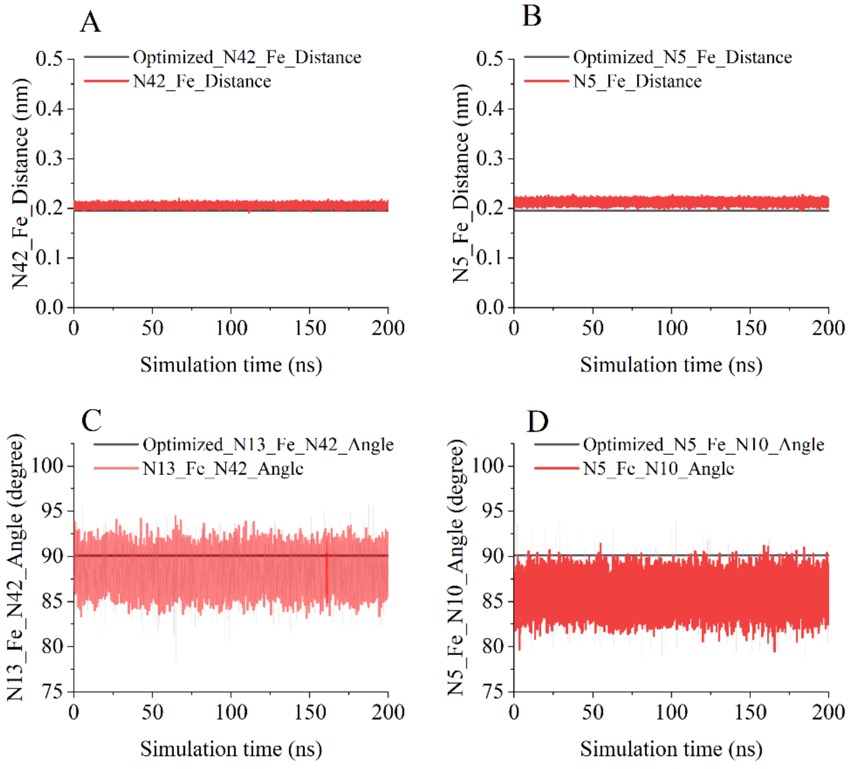

**Supplementary Figure 2.** Comparison between the selected angles and distances during the MD simulation and the optimised geometry distances and angles.

### Validation of the AlphaFold predicted structure of VP_fold and MnP_fold

The structures of VP_fold and MnP_fold were predicted using AlphFold3, and the prediction was validated before proceeding to the molecular dynamics simulation. To validate the predicted structure, the backbone atoms RMSD was calculated for the VP_fold and MnP_fold structures. The resulting RMSD of 0.8 and 1.19, respectively, aligned with VP (PDB id: 2boq), indicating good prediction results.

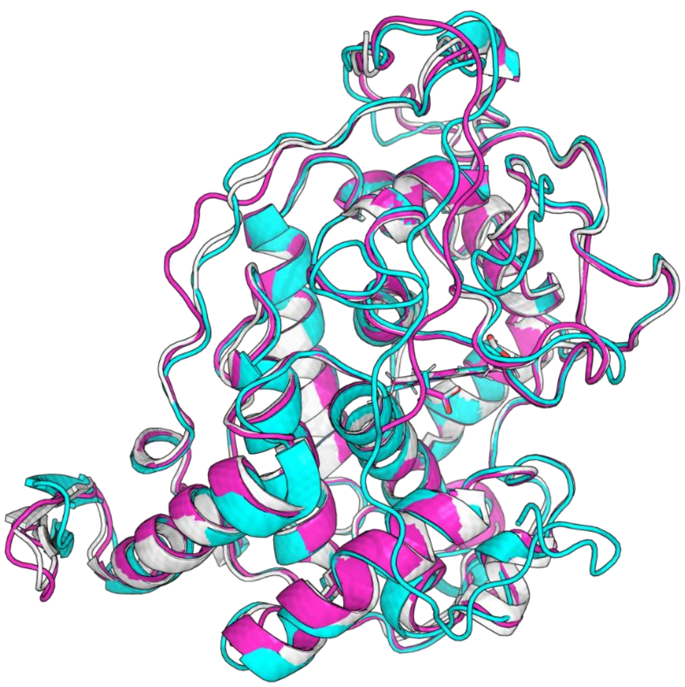

**Supplementary Figure 3.** Structural alignment of VP_fold and MnP_fold to the crystal structure of VP (PDB id 2boq). The alignments results indicate an agreement for general fold, especially for the helical positions, indicating a good prediction result.

### Purification of VP and MnP.

Both VP (Supplementary Figure 4a) and MnP (result not shown) were subjected to anion-exchange chromatography and only fractions corresponding to the 407 nm heme peak and VP and MnP activities were pooled. The SDS-PAGE showed a closely-spaced doublet band for VP (Supplementary Figure 4, inset) as described previously (Ertan et al. 2012) and a single band for MnP (Supplementary Figure 4b), implying purification to homogeneity level for both proteins.

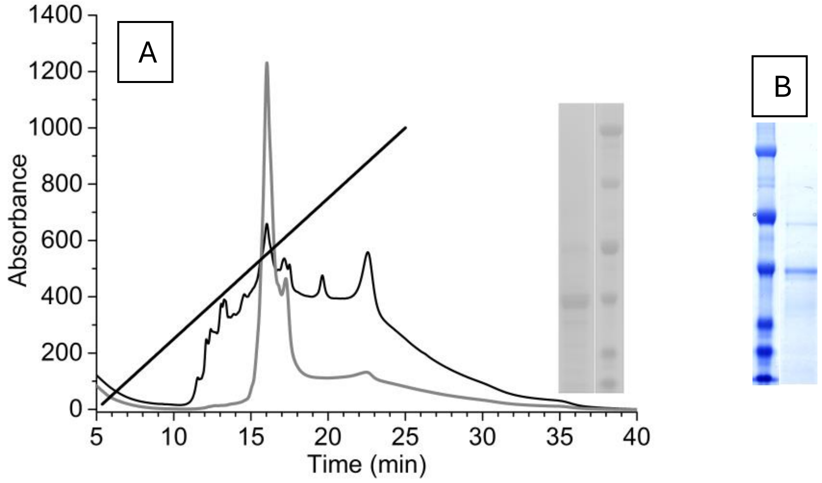

**Supplementary Figure 4**. **A**: Chromatographic purification and electrophoretic analysis of VP. Chromatogram showing anion-exchange column chromatography of VP. Dark line = $A_{280,}$ grey line = $A_{407.}$ The diagonal line is the NaCl gradient from 0–1 M. **Inset:** 12 % SDS-PAGE showing purified VP (left lane) and molecular weight markers (right lane). Makers size from top to bottom: β-galactosidase (120 kDa), phosphorylase B (97 kDa), bovine serum albumin (67 kDa), ovalbumin (45 kDa), carbonic anhydrase (30 kDa), lysozyme (14.4 kDa). **B:** 12 % SDS-PAGE of MnP after purification on anion-exchange chromatography as described for VP. Right lane: purified MnP. Left lane: molecular weight markers. Makers size from top to bottom: phosphorylase B (97 kDa), bovine serum albumin (67 kDa), ovalbumin (45 kDa), carbonic anhydrase (30 kDa), myoglobin (18 kDa), lysozyme (14.4 kDa).
